# Biological upgrading of C1-C2 products of electrocatalytic CO_2_ reduction to C4-C6 carboxylates

**DOI:** 10.64898/2026.08.03.741547

**Authors:** Chao Xu, Jonathan K. Otten, John D. Hill, Noah B. Wills, Eleftherios T. Papoutsakis

## Abstract

**Background:** Microbial chain-elongation by *Clostridium kluyveri* using the products (acetate and ethanol) derived from the electrocatalytic CO_2_ reduction reaction (CO_2_RR) represents a unique sustainable strategy for producing C4-C6 chemicals from CO_2_. However, direct integration of electrocatalytic effluents with anaerobic bioprocesses is often impeded by the physiological incompatibility between electrocatalytic product streams and microbial metabolism. Specifically, CO_2_RR effluents commonly contain formate, which cannot be utilized by *C. kluyveri* for chain elongation and therefore reduces the overall carbon efficiency of CO_2_ conversion to C4–C6 chemicals. Moreover, both formate and the elevated phosphate concentrations typical of electrochemical reaction solutions may inhibit microbial growth.

**Results:** We show that formate at concentrations of up to 50 mM did not inhibit the growth of or the chain elongation by *C. kluyveri*. Based on this finding, we developed a modular two-step bioprocess. In the first step, the acetogen *Clostridium ljungdahlii* converts formate in CO_2_RR product mixtures into acetate, thereby generating additional substrates for second-step *C. kluyveri*-driven chain elongation, thus increasing the CO_2_RR carbon-conversion efficiency to C– C6 chemicals. To address the issue of *C. ljungdahlii*’s inhibition by high phosphate concentrations in electrocatalytic solutions, we explored the use of *C. ljungdahlii* biofilms for the first, i.e. the formate-conversion, step. *C. ljungdahlii* biofilms exhibit tolerance to concentrated electrolytes, enabling the conversion of up to 50 mM formate in CO_2_RR solutions.

**Conclusions:** The demonstrated two-step process constitutes the basis for the development of a robust and carbon-efficient biological process for the scalable upgrading of C1–C2 CO_2_RR products into higher-value C4–C6 chemicals.

## INTRODUCTION

Increasing interest in carbon-neutral chemical manufacturing has spurred the development of integrated electrosynthesis–bioprocessing technologies that can convert carbon dioxide (CO) into fuels (1), plastics (2), and food (3). Recent advances in electrocatalytic CO reduction reactions (CO RR) enable the selective generation of C1–C3 products such as formate, acetate, ethanol, and propanol using electricity (4, 5). Despite these advances, direct commercialization of short-chain CO RR products is often constrained by their relatively low energy density and limited market value of C1-C3 products (6). Consequently, biologically upgrading CO RR-derived intermediates into higher-value C4–C8 molecules has emerged as a compelling route to improve carbon valorization and process economics.

Among anaerobic upgrading pathways, reverse β-oxidation–based chain elongation is particularly attractive because it efficiently converts low-carbon chain precursors (notably ethanol and acetate) into medium-chain fatty acids (MCFAs)(7, 8). *Clostridium kluyveri* (which does not utilize sugars) is the best-characterized chain-elongating organism, converting ethanol and acetate into butyrate and hexanoate (and also octanoate) via thermodynamically favorable reactions (9, 10). This makes *C. kluyveri* an appealing biocatalyst for upgrading CO RR derived C1-C2 chemicals. Of note, CO as co-substrate is critical in *C. kluyveri’s* physiology, as has been recognized by the pioneering studies of Barker and coworkers. Notably, Tomlinson and Barker (11) demonstrated that more than 25% of cell carbon originates from CO . Recent bioreactor studies have confirmed a direct relationship between CO availability, growth, and chain elongation performance (12). However, coupling *C. kluyveri* directly with CO RR outputs faces a major compatibility challenge: CO RR product mixtures frequently and typically contain formate (see Additional file 1), at concentrations that increase with longer operation of the electrocatalyzer aiming to increase the concentrations of ethanol and acetate. Formate is not a substrate for chain elongation and is toxic to most microbial cells including some *Clostridium* species (13). The impact of formate on *C. kluyveri* growth, physiology, and chain-elongation performance has not been systematically evaluated. In addition, CO RR electrolytes commonly employ concentrated phosphate buffers, resulting in elevated ionic strengths and osmotic pressures that can further compromise *C. kluyveri* viability and metabolic activity. Indeed, potassium phosphate buffers in the 100–200 mM concentration range have been widely employed in studies of aqueous CO electroreduction because of their relatively high buffering capacity and ability to mitigate bulk pH excursions (14). Together, formate co-production and high-phosphate conditions represent key barriers to seamlessly integrating CO RR with microbial chain elongation.

There are few organisms that in the absence of sugars or oxygen can use formate as a substrate for growth or convert it to non-toxic chemicals that can be further metabolized by *C. kluyveri*, notably ethanol, acetate, and CO_2_. In fact, only acetogens meet this criterion. Thus, a rational strategy to overcome the aforementioned barriers of chain elongation is to incorporate an acetogenic partner that can consume formate and simultaneously supply chain-elongation substrates, notably acetate, CO_2_ (which enhances chain elongation by *C. kluyveri*) and possibly ethanol under proper culture conditions. Several clostridial acetogens, including *Clostridium ljungdahlii* and *Acetobacterium woodii*, convert formate to acetate via the Wood–Ljungdahl pathway (WLP)(15–17). In a co-culture context, such acetogens can function as a metabolic “formate sink” while generating acetate and CO_2_ (depending on conditions) ethanol, thereby providing the key precursors required by *C. kluyveri* for chain elongation. Moreover, phylogenetic relatedness within the clostridial lineage may facilitate interspecies compatibility and stable metabolite exchange (10, 17, 18). Consistent with this concept, we and others have observed robust *C. ljungdahlii–C. kluyveri* cocultures that support coordinated growth and sustained chain elongation (10, 18, 19), suggesting that *C. ljungdahlii* is a promising partner for converting electrosynthetic produced formate to C4-C8 chemicals via chain elongation.

From a bioprocessing perspective, realizing this coupling in scalable and continuous reactors requires robust strategies to retain and reuse the acetogenic partner while maintaining its formate conversion function. Immobilizing cells through biofilm formation is a well-established route to enhance biomass retention, operational stability, and catalyst reusability in continuous systems (20–22). Accordingly, *C. ljungdahlii* biofilms present an attractive engineering solution for sustained formate conversion in reactor configurations such as packed-bed systems. Philips et al. reported NaCl stress–induced biofilm formation by *C. ljungdahlii* and used transcriptome analysis to probe underlying mechanisms of biofilm formation, demonstrating cell attachment and biofilm formation on multiple surfaces (e.g., glass, graphite, and glassy carbon)(23). However, cellular viability and metabolic activity in developing *C. ljungdahlii* biofilms were not evaluated, and the potential of *C. ljungdahlii* biofilms for bioprocessing applications was not explored.

Here, we develop a two-stage, potentially scaleable bioprocess in which CO electroreduction– derived C1–C2 products are upgraded into C4–C6 chemicals, with a focus on butyrate and hexanoate production using, sequentially, *C. ljungdahlii* and *C. kluyveri*. In this framework, *C. ljungdahlii* converts formate into a chain-elongation–compatible product spectrum (acetate and ethanol), while *C. kluyveri* elongates these intermediates to MCFAs. We also examined this biological system using high phosphate concentrations (up to 140 mM total phosphate), which fall well within the range commonly utilized in fundamental CO RR investigations as cited above. In parallel, we evaluated *C. ljungdahlii* biofilms formed on metal and glass supports as a formate-converting biocatalyst, highlighting their potential as packing media for packed-bed reactor operation. Collectively, this work establishes a practical route to upgrade CO electroreduction outputs via anaerobic bioprocessing and provides a conceptual and engineering framework for electro-bio hybrid platforms targeting renewable C4–C8 platform chemicals.

## METHODS

### Strains and Media

Wild-type *Clostridium ljungdahlii* (DSM 13528) and wild-type *Clostridium kluyveri* (ATCC 8527) were cultivated either as monocultures or in sequential co-cultures using media with different carbon sources and buffer concentrations based on Turbo CGM (TCGM) (24) medium, as summarized in Table 1.

**Table 1.** Media used in this study.

| Organism/Culture | Medium | Experimental Purpose | Carbon Source(s) | pH | KH<sub>2</sub>PO<sub>4</sub> (mM) | K<sub>2</sub>HPO<sub>4</sub> (mM) | Other Components |
| --- | --- | --- | --- | --- | --- | --- | --- |
| *C. ljungdahlii* monoculture | Clj-TCGM-Standard | In Section 2.2 | 27.8 mM fructose | 6.8 | 7.3 | 7.2 | — |
|  | Clj-TCGM-Control | In Section 2.4 | 30 mM fructose | 6.8 | 7.3 | 7.2 | — |
|  | Clj-TCGM-Formate-Growth |  | 50 mM or 100 mM sodium formate | 6.8 | 7.3 | 7.2 | — |
|  | Clj-TCGM-High-Buffer | In Section 2.5 | 27.8 mM fructose | 6.8 | 44.1 | 95.4 | — |
|  | Clj-TCGM-Biofilm | In Section 2.9 | 55.5 mM fructose | 6.8 | 7.3 | 7.2 | 200 mM sodium chloride |
|  | Clj-TCGM-Formate-Conversion | In Section 2.11 | 30 mM, 50 mM, or 70 mM sodium formate | 6.8 | 44.1 | 95.4 | — |
|  | Clj-TCGM-Formate-Conversion in mixture |  | 50 mM or 70 mM sodium formate in the presence of 40 mM sodium acetate and 100 mM ethanol | 6.8 | 44.1 | 95.4 | — |
| *C. kluyveri* monoculture | Ckl-TCGM-Standard | In Section 2.2 | 97.5 mM sodium acetate, 343 mM ethanol | 6.8 | 7.3 | 7.2 | 30 mM sodium bicarbonate, 1.7 mM L-cysteine HCl |
|  | Ckl-TCGM-Formate | In Section 2.3 | 20 mM or 50 mM sodium formate, 40 mM sodium acetate, 100 mM ethanol | 6.8 | 7.3 | 7.2 | — |
|  | Ckl-TCGM-High-Buffer | In Section 2.5 | 97.5 mM sodium acetate, 343 mM ethanol | 6.8 | 44.1 | 95.4 | 30 mM sodium bicarbonate, 1.7 mM L-cysteine HCl |
| *C. ljungdahlii* and *C. kluyveri* coculture | Coculture-TCGM-Standard | In Section 2.5 | 50 mM sodium formate, 40 mM sodium acetate, | 6.8 | 7.3 | 7.2 | — |

| Organism/Culture | Medium | Experimental Purpose | Carbon Source(s) | pH | $\text{KH}_2\text{PO}_4$ (mM) | $\text{K}_2\text{HPO}_4$ (mM) | Other Components |
| --- | --- | --- | --- | --- | --- | --- | --- |
|  |  |  | 100 mM ethanol |  |  |  |  |
|  | Coculture-TCGM-High-Buffer | In Section 2.7 | 50 mM sodium formate, 40 mM sodium acetate, 100 mM ethanol | 6.8 | 44.1 | 95.4 | — |
|  | Coculture-TCGM-High-Buffer (80 mM acetate and 200 mM ethanol) | In Section 2.8 | 50 mM sodium formate, 80 mM sodium acetate, 200 mM ethanol | 6.8 | 44.1 | 95.4 | — |
|  | Coculture-TCGM-High-Buffer (120 mM acetate and 300 mM ethanol) |  | 50 mM sodium formate, 120 mM sodium acetate, 300 mM ethanol | 6.8 | 44.1 | 95.4 | — |

### Preparation of Seed Cultures for *C. ljungdahlii* and *C. kluyveri*

*C. kluyveri* was revived from frozen stocks in 30 mL of Ckl-TCGM-Standard medium (Table 1) in 160 mL serum bottles. *C. ljungdahlii* was grown under a 20-psi headspace of 80/20 (H /CO, v/v). *C. kluyveri* was grown under the atmosphere of the gas (5% H, 10% CO, 85% N) in the anaerobic glove-box (Forma). *C. ljungdahlii* cultures was incubated at 37 °C and 90 rpm on a rotary shaker, whereas *C. kluyveri* cultures were incubated at 37 °C in a small incubator inside the glove-box. OD and pH of both *C. ljungdahlii* and *C. kluyveri* were monitored periodically, and cultures at exponential phase (OD = 0.5–1.0) were used as seed cultures.

### Examination of the Impact of Formate on *C. kluyveri* Cultures

*C. kluyveri* seed cultures were inoculated at 20% (v/v) into 30 mL Ckl-TCGM-Formate medium (Table 1) containing either 0 (control), 20 or 50 mM sodium formate in 160 mL serum bottles. All handling was performed in an anaerobic glove box (5% H, 10% CO, 85% N). Cultures were incubated at 37 °C and 90 rpm, and OD and pH were measured every 12 h.

### *C. ljungdahlii* Growth on Formate

*C. ljungdahlii* seed cultures were inoculated at 20% (v/v) into 30 mL Clj-TCGM-Formate-Growth medium (Table 1) containing either 0 (control), 50 or 100 mM sodium formate in 160 mL serum bottles. After inoculation, bottle headspaces were flushed with 80/20 (v/v) H /CO at 20 psi for 2 min. Cultures were incubated at 37 °C and 90 rpm, and OD, pH, and metabolite concentrations were measured every 12 h.

### Cultures of *C. ljungdahlii* and *C. kluyveri* Under High-Phosphate Buffer Concentrations

For *C. ljungdahlii*, seed cultures were inoculated (20%, v/v) into 30 mL of Clj-TCGM-High-Buffer medium (Table 1) in 160 mL serum bottles. Cultures grown in Clj-TCGM-Standard medium served as controls. After inoculation, the headspace was flushed with 80/20 (H /CO, v/v) at 20 psi for 2 min. Cultures were incubated at 37 °C and 90 rpm, and OD and pH were measured every 24 h.

For *C. kluyveri*, seed cultures were inoculated (20%, v/v) into 30 mL of Ckl-TCGM-High-Buffer medium in 160 mL serum bottles. Cultures grown in Ckl-TCGM-Standard medium served as controls. All procedures were performed in the anaerobic glove box; bottles were sealed and incubated at 37 °C and 90 rpm. OD and pH were measured every 24 h.

### Two-Step Sequential Cultures of *C. ljungdahlii* and *C. kluyveri*

Two-step cultures were performed sequentially as follows.

### Step 1: Formate Conversion by *C. ljungdahlii*

*C. ljungdahlii* cells were harvested by centrifugation (4000 rpm, 4 °C, 10 min) and resuspended in 1 mL of Clj-TCGM-Standard medium (Table 1). The suspension was inoculated into 60 mL of Coculture-TCGM-Standard medium in 250 mL serum bottles to achieve an initial OD of 0.5. Cultures were incubated under an N headspace at 37 °C and 90 rpm until formate was fully consumed. After formate depletion was confirmed, the culture pH was adjusted to 6.8–7.0.

### Step 2: Chain Elongation by *C. kluyveri*

*C. kluyveri* cells were harvested by centrifugation (4000 rpm, 4 °C, 10 min), resuspended in 1 mL of Ckl-TCGM-Standard medium (Table 1), and inoculated into serum bottles containing the *C. ljungdahlii*–treated (formate-depleted) medium. Bottles were sealed in the anaerobic glove box without additional gas flushing and then transferred to be incubated at 37 °C and 90 rpm. OD, pH, and metabolite concentrations were monitored every 24 h to quantify growth and medium-chain fatty acid production.

### Two-Step Sequential Cultures of *C. ljungdahlii* and *C. kluyveri* Under High-Phosphate Buffer Conditions at Different Inoculation Densities

Two-step cultures under high-phosphate buffer conditions were performed as described above, except that Coculture-TCGM-High-Buffer medium (Table 1) was used and the initial OD for each stage was varied. Specifically, *C. ljungdahlii* and *C. kluyveri* were inoculated at initial OD values of 0.1, 0.3, or 0.5 for the formate-conversion and chain-elongation stages, respectively. All other conditions (centrifugation, incubation conditions, pH adjustment, and monitoring frequency) were identical to those described above

### Two-Step Sequential Cultures of *C. ljungdahlii* and *C. kluyveri* Under High-Phosphate Buffer Conditions at Elevated Acetate and Ethanol Concentrations

Two-step chain cultures were performed as described above using Coculture-TCGM-High-Buffer medium, except that acetate and ethanol concentrations were increased while the initial OD was fixed at 0.5 for each stage. For the formate-conversion stage, Coculture-TCGM-High-Buffer medium contained either 50 mM formate, 80 mM acetate, and 200 mM ethanol or 50 mM formate, 120 mM acetate, and 300 mM ethanol. After completion of formate conversion and adjustment of pH to 6.8–7.0, *C. kluyveri* was inoculated to initiate chain elongation. All remaining procedures and monitoring were performed as described above.

### Biofilm Formation of *C. ljungdahlii* on Saddle-Shaped Metal Packing and Glass Surfaces

Seed cultures of *C. ljungdahlii* were inoculated at 20% (v/v) into Clj-TCGM-Biofilm medium. The headspace was flushed with 80/20 (H /CO, v/v) at 20 psi for 2 min, and cultures were incubated at 37 °C and 90 rpm. OD and pH were monitored until OD reached 2.0 to obtain biofilm inoculum (“biofilm culture”) for surface colonization.

### Saddle-Shaped Metal Packing

Forty pieces of saddle-shaped metal packing (total mass ≈ 690 mg) were placed into sterile Hungate tubes containing 15 mL of the biofilm culture and incubated statically for 3 days at 37 with the glove box gas atmosphere. Saddle-shaped (half-cylinders with two corners bent slightly outward) stainless-steel distillation-column packing (0.16”x0.16”), a Cannon Instrument Company product was purchased from Ace Glass Inc., NJ, US (Cat. No. 6624-04). The supernatant was removed, and saddles were gently washed three times with sterile, deoxygenated PBS (pH 7.4) to remove non-adherent cells while minimizing biofilm detachment.

### Glass Surfaces

Biofilms on flat glass surfaces to enable better imaging were developed on the glass bottom of a cell culture chamber slide with 8 wells (1 cm^2^ bottom surface area)) suitable for microscopic examination (Cat. No. 80806, Ibidi GmbH, Gräfelfing, Germany). 0.5 mL of the biofilm culture was added to each chamber (although 300 μl volume per well is cited in the product specs) of and incubated statically for 3 days. The supernatant was removed and each chamber was gently washed three times with sterile PBS (pH 7.4) to remove unattached planktonic cells while minimizing disruption of surface-attached biofilms

### Characterization of *C. ljungdahlii* Biofilms

#### Crystal Violet Staining Assay for Quantifying *C. ljungdahlii* Biofilm Biomass on Metal Packing

Biofilm biomass on saddle-shaped metal packing was quantified using a crystal violet (CV) assay (25). Saddle-supported biofilms were prepared as described above and washed three times with sterile PBS (pH 7.4). Washed saddles (40 pieces per tube) were stained with 2 mL of 0.1% (w/v) crystal violet for 15 min at room temperature. Excess stain was removed, and saddles were rinsed 3–5 times with 5 mL PBS or deionized water until rinsed solution was visibly clear. Saddles were air-dried at room temperature. Bound CV was solubilized with 5 mL of 30% (v/v) acetic acid for 15 min at room temperature with gentle mixing. The destaining solution was collected, briefly vortexed, and absorbance was measured at 570 nm (OD) using 30% (v/v) acetic acid as the blank. Biofilm biomass was reported as OD per tube (40 saddles per tube). At least three biological replicates were performed.

### Scanning Electron Microscopy (SEM) Analysis of Saddle-Supported Biofilms

Scanning electron microscopy (SEM) was used to examine cell attachment and surface coverage of saddle-supported *C. ljungdahlii* biofilms. Samples were fixed, dehydrated through graded ethanol, and dried as reported (25). Dried samples were sputter-coated with a 2.5-nm Au–Pt layer using a Leica EM ACE600 high-vacuum coater (Leica Microsystems, Wetzlar, Germany). SEM imaging was performed using a Thermo Scientific Apreo VolumeScope scanning electron microscope (Thermo Fisher Scientific, Hillsboro, OR, USA) at an accelerating voltage of 5–10 kV.

### Confocal Laser Scanning Microscopy (CLSM) Analysis of Biofilm Thickness and Viability Based on RNA FISH

Confocal laser scanning microscopy (CLSM) was used to characterize the three-dimensional architecture and viable-cell distribution of *C. ljungdahlii* biofilms. Biofilms were gently rinsed with sterile PBS (pH 7.4) to remove unattached cells. Viability was assessed using an rRNA-targeted fluorescence in situ hybridization (RNA-FISH) workflow previously established in our laboratory (18). Confocal imaging was performed using an ANDOR Dragonfly spinning-disk confocal microscope (Oxford Instruments Andor) (18).

### *C. ljungdahlii* Biofilms for Formate Conversion

After biofilm formation on metal saddles, spent medium was removed and saddles were gently rinsed with sterile PBS (pH 7.4) to remove unattached cells. Biofilm-laden saddles were transferred using tweezers into fresh Clj-TCGM-Formate-Conversion medium supplemented with sodium formate (30, 50, or 70 mM) for formate-conversion assays. To evaluate electrolysis-relevant mixed-substrate conditions, biofilm-laden saddles were also transferred into Formate-Conversion medium supplemented with 40 mM sodium acetate, 100 mM ethanol, and either 50 or 70 mM sodium formate (Clj-TCGM-Formate-Conversion in mixture, as specified for the medium of each experiment). Headspaces were set to either 100% N or 80/20 (H /CO, v/v) to assess the effect of gas composition. All incubations were conducted in sealed Hungate tubes under anaerobic conditions.

### Detection of Metabolites

Metabolites in the fermentation broth were quantified using high-performance liquid chromatography (HPLC). Analyses were carried out on an Agilent system equipped with a Bio-Rad Aminex HPX-87H separation column. The column was operated with 5 mM sulfuric acid as the mobile phase at a flow rate of 0.5 mL/min (26).

## RESULTS AND DISCUSSION

### Formate Does Not Impair the Growth or Chain-Elongation Performance of *C. kluyveri*

CO electrocatalytic effluents typically contain formate, acetate, and ethanol (see Additional file 1). However, *C. kluyveri* can use only acetate and ethanol as substrates for chain elongation. Also, prior studies have suggested that even small concentrations of formate can trigger an “acid crash” in certain *Clostridium* species (13). This has raised concerns regarding the direct use of CO RR products in *C. kluyveri* cultures. To evaluate the effect of formate on *C. kluyveri*, we cultured *C. kluyveri* in media supplemented with 20 mM or 50 mM formate. As shown in Fig. 1A-C, no obvious differences were observed between the control and formate-supplemented cultures. Following inoculation, the OD_600_ increased while the pH decreased rapidly (indicative of acid uptake), indicating active growth by *C. kluyveri*. After 24 h, the OD_600_ began to decline and the pH became relatively stable, likely due to the depletion of acetate or ethanol. Metabolite analysis (Fig. 1D) showed that *C. kluyveri* did not consume formate at either 20 or 50 mM, while the production of butyrate and hexanoate remained comparable to that in the control. These results indicate that the presence of formate did not alter the metabolic pattern of *C. kluyveri* during chain elongation. To our knowledge, this study provides the first evidence that formate at concentrations of up to 50 mM does not inhibit *C. kluyveri* cultures in the presence of sufficient nutrients, that is, ethanol and acetate.

**Figure 1.**
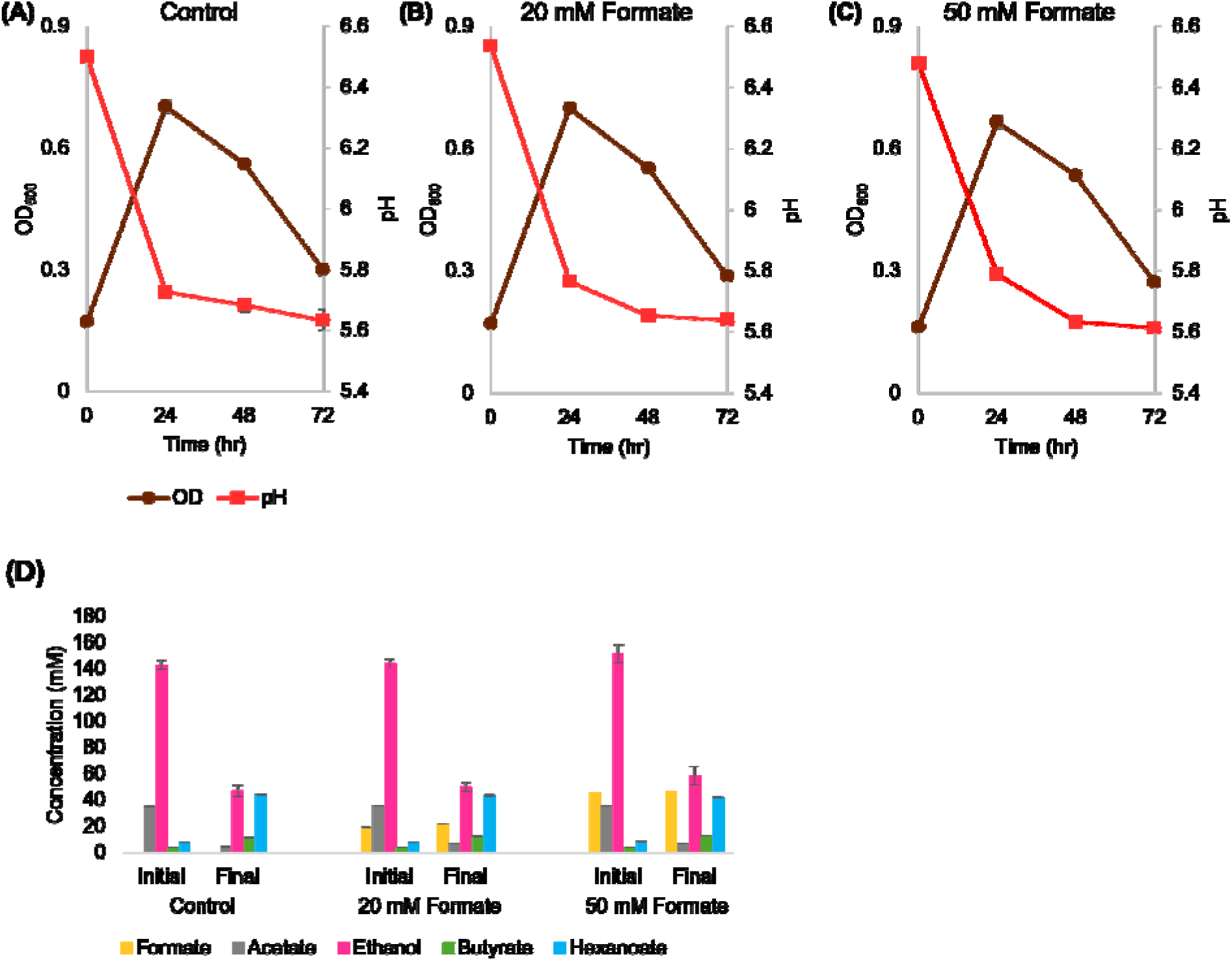
Biomass formation kinetics (in OD_600_) and pH profiles of *C. kluyveri* grown with 0 mM formate (A), 20 mM formate (B), and 50 mM formate (C). (D) The initial and final product and substrate concentrations (mM). (n=3).

### Physiological Window for Formate Utilization by *C. ljungdahlii*

As anacetogen, *C. ljungdahlii* encodes the native Wood–Ljungdahl pathway (WLP), which in principle enables assimilation of formate into acetyl-CoA and downstream acetate (27). This metabolic capability makes *C. ljungdahlii* a promising organism for utilizing formate in formate-rich CO RR products. To define the physiological limits of formate utilization in planktonic cultures, *C. ljungdahlii* was cultivated under an 80:20 (v/v) H /CO atmosphere (typically used for autotrophic cultures of this and other acetogens) with either 30 mM fructose (control) or sodium formate as the sole soluble carbon source (50 or 100 mM) in addition to the CO_2_ available in the headspace.

As shown in Fig. 2A, fructose-supported cultures grew rapidly and acidified the medium, accompanied by conversion of fructose to acetate, consistent with canonical acetogenic growth. In contrast, when formate was provided as the sole carbon source, growth was markedly slower and pH shifts were attenuated. Importantly, as shown in Fig. 2B, 50 mM formate was completely consumed within 48 h with concomitant acetate formation, confirming that exogenous formate can be assimilated through the WLP under the tested conditions. However, increasing the initial formate concentration to 100 mM fully suppressed growth and acetate production, indicating a concentration-dependent toxicity threshold that constrains the operational window for formate-based bioconversion in planktonic *C. ljungdahlii*. The tolerance observed here is higher than that reported by Ramió-Pujol and Ganigué (28), who observed complete inhibition of *C. ljungdahlii* at sodium formate concentrations as low as 55 mM with a starting pH of 6.0 (similar to our starting condition). Also, we observed complete consumption of 50 mM of formate in 48 hours, where Ramió-Pujol and Ganigué (28) required 7 days to consume 25 mM of formate. In our experiments, the wild-type strain retained clear metabolic activity at 50 mM of formate.

**Figure 2.**
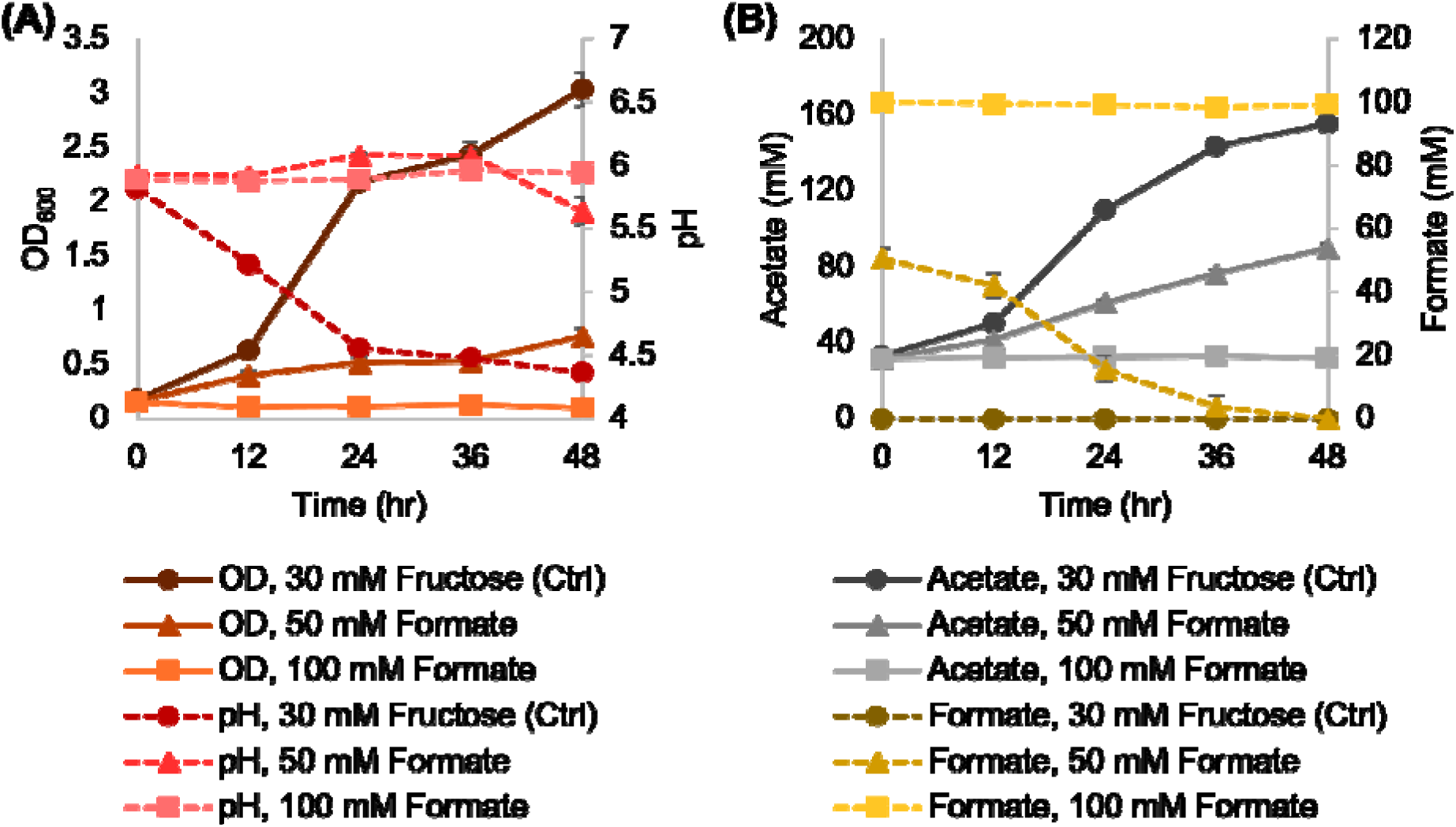
(A) Biomass formation kinetics (in OD_600_) and pH profiles of *C. ljungdahlii* grown with 0 mM formate and 30 mM fructose, 50 mM formate, and 100 mM formate. (B) Acetate and formate concentration over time. (n=2-3).

### Two-Step Chain Elongation in a Sequential *C. ljungdahlii–C. kluyveri* Coculture

Electrochemical CO reduction typically yields a mixed product stream containing formate together with acetate and ethanol. Direct upgrading of such effluents via *C. kluyveri-*catalyzed chain elongation may result in the waste of formate, since it cannot be utilized by *C. kluyveri*, thereby lowering the carbon efficiency of the whole process. To circumvent this bottleneck, we implemented a sequential two-stage biological interface. In the 1^st^ stage, *C. ljungdahlii* serves as a selective formate-conversion agent by assimilating formate via the WLP and converting it primarily into acetate, thereby transforming an electrochemical byproduct into substrates compatible with chain elongation. To emulate a CO electrolysis-like effluent, cultures were initially supplied with 50 mM formate, 40 mM acetate, and 100 mM ethanol.

During the formate-conversion stage (Fig. 3a), *C. ljungdahlii* showed limited biomass accumulation yet maintained measurable metabolic activity. Consistent with this, formate was selectively consumed with a corresponding increase in acetate (by an average of 10 mM) and in ethanol formation (by an average of 6.5 mM) by the end of the 1^st^ stage (Fig. 3b) upon the consumption of an average of 45 mM formate, with about 7.5 mM formate left in the medium These data demonstrate efficient biological formate conversion with a good stoichiometric conversion of formate: with one mole of either acetate or ethanol deriving from two moles of formate, carbon recovery for this set of experiments (16.5 2=33 mM, the balance of 45 33 = 12 mM formate is assumed to be oxidized to CO_2_). Upon formate depletion, inoculation with *C. kluyveri* triggered rapid biomass accumulation (Fig. 3a) accompanied by accelerated consumption of acetate and ethanol (Fig. 3b), resulting in production of 17.4 ± 0.2 mM butyrate and 40.5 ± 0.3 mM hexanoate. These results demonstrate that biological conversion of formate by *C. ljungdahlii* enables effective upgrading of electrolysis-derived substrates through *C. kluyveri* chain elongation.

**Figure 3.**
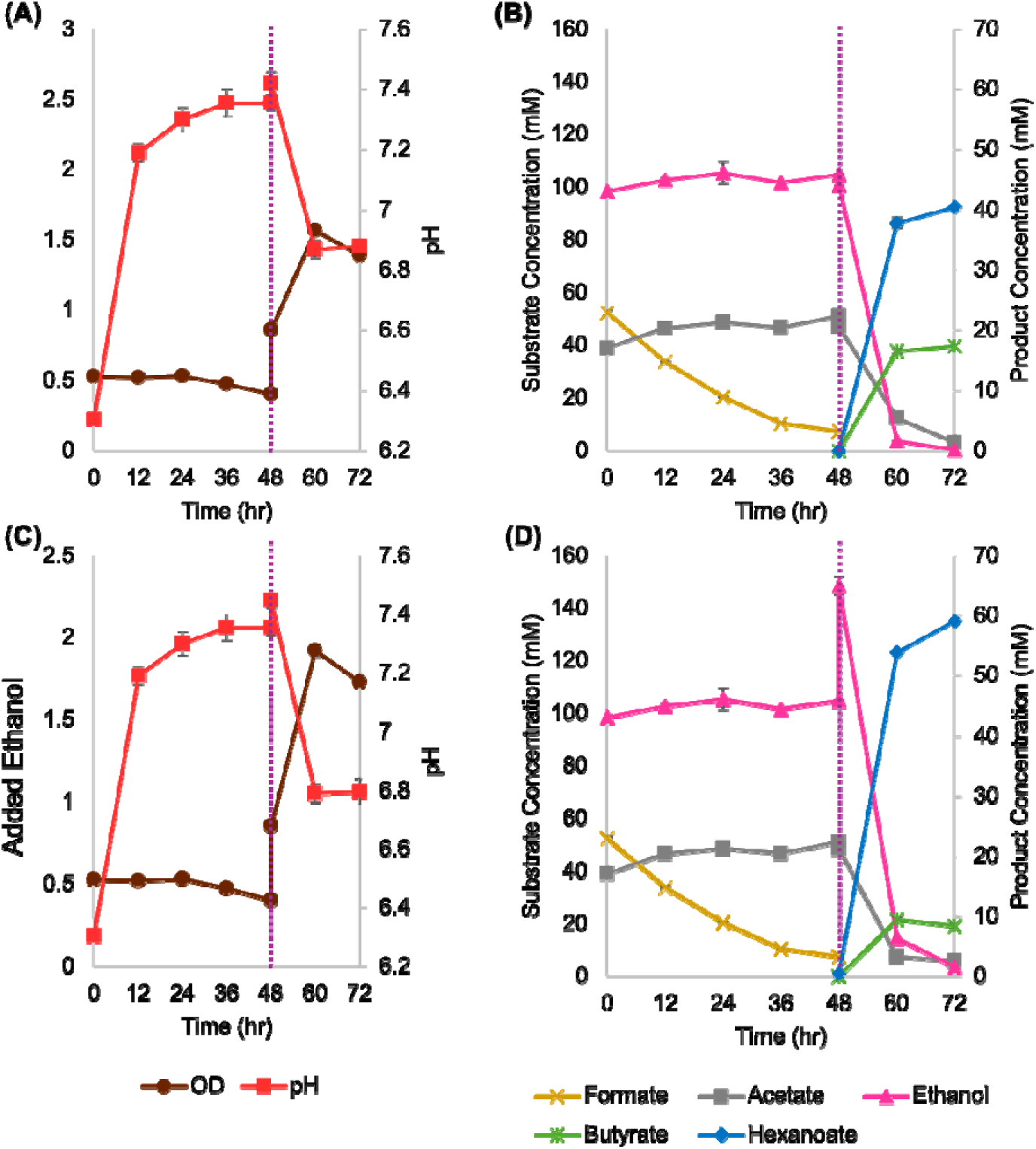
Biomass formation kinetics (in OD_600_), pH profiles, and substrate consumption and product formation kinetics of sequential cocultures of *C. ljungdahlii* and *C. kluyveri*. The dashed vertical lines represent the inoculation time of *C. kluyveri*. C and D: 50 mM additional ethanol was supplemented at 48 hours. Two biological replicates were employed per condition.

To further evaluate system versatility under increased electron and carbon availability, we increased the ethanol load during the 2^nd^ stage. After formate depletion by *C. ljungdahlii* but prior to *C. kluyveri* inoculation, an additional 50 mM ethanol was supplemented, raising the total ethanol concentration to 150 mM. Based on previous research in our lab demonstrating chain elongation by *C. kluyveri*, it is important to have ethanol-to-acetate ratio to be 3:1 (10). With this ethanol addition, biomass accumulation increased relative to the control (Fig. 3c), and acetate and ethanol were depleted within 24 hours (Fig. 3d), yielding 8.4 ± 0.6 mM butyrate and 59.1 ± 0.3 mM hexanoate. Thus, with increased substrate loading, formate, ethanol and acetate input can be effectively converted to MCFAs.

Collectively, these results demonstrate functional metabolic integration in the two-stage platform: *C. ljungdahlii* converts formate into ethanol (conserving carbon and reducing equivalents), whereas *C. kluyveri* subsequently upgrades acetate/ethanol into higher-value MCFAs.

### Testing the Ability of *C. ljungdahlii* and *C. kluyveri* to Grow Under High Concentrations of Phosphate Buffers Used in Electrocatalytic Conversion of CO_2_

Bridging CO electroreduction with downstream fermentation requires compatibility between electrolysis-relevant electrolyte compositions and microbial physiology. High-strength phosphate buffers (in the range of 100 - 500 mM phosphate concentrations)(14) are typically used in CO RR to stabilize pH and improve conductivity; however, their effects on acetogenic and chain-elongating systems remain insufficiently characterized. We therefore assessed the growth and metabolic responses of *C. ljungdahlii* and *C. kluyveri* under elevated phosphate conditions (a total of about 140 mM, Table 1) representative of electrolysis operation.

As shown in Fig. 4A, *C*. ljungdahlii displayed reduced growth kinetics relative to low phosphate controls (about 15 mM, Table 1) but remained viable and metabolically active throughout the culture (as evidenced by strong pH drop). In contrast, in Fig. 4B, *C*. kluyveri showed minimal sensitivity, with growth behavior and biomass yield largely unchanged under high phosphate-buffer concentrations. In addition to physiological tolerance, the high buffer concentration provided an operational advantage by suppressing large pH drops observed in low-buffer controls, thereby reducing acid-stress–associated inhibition.

**Figure 4.**
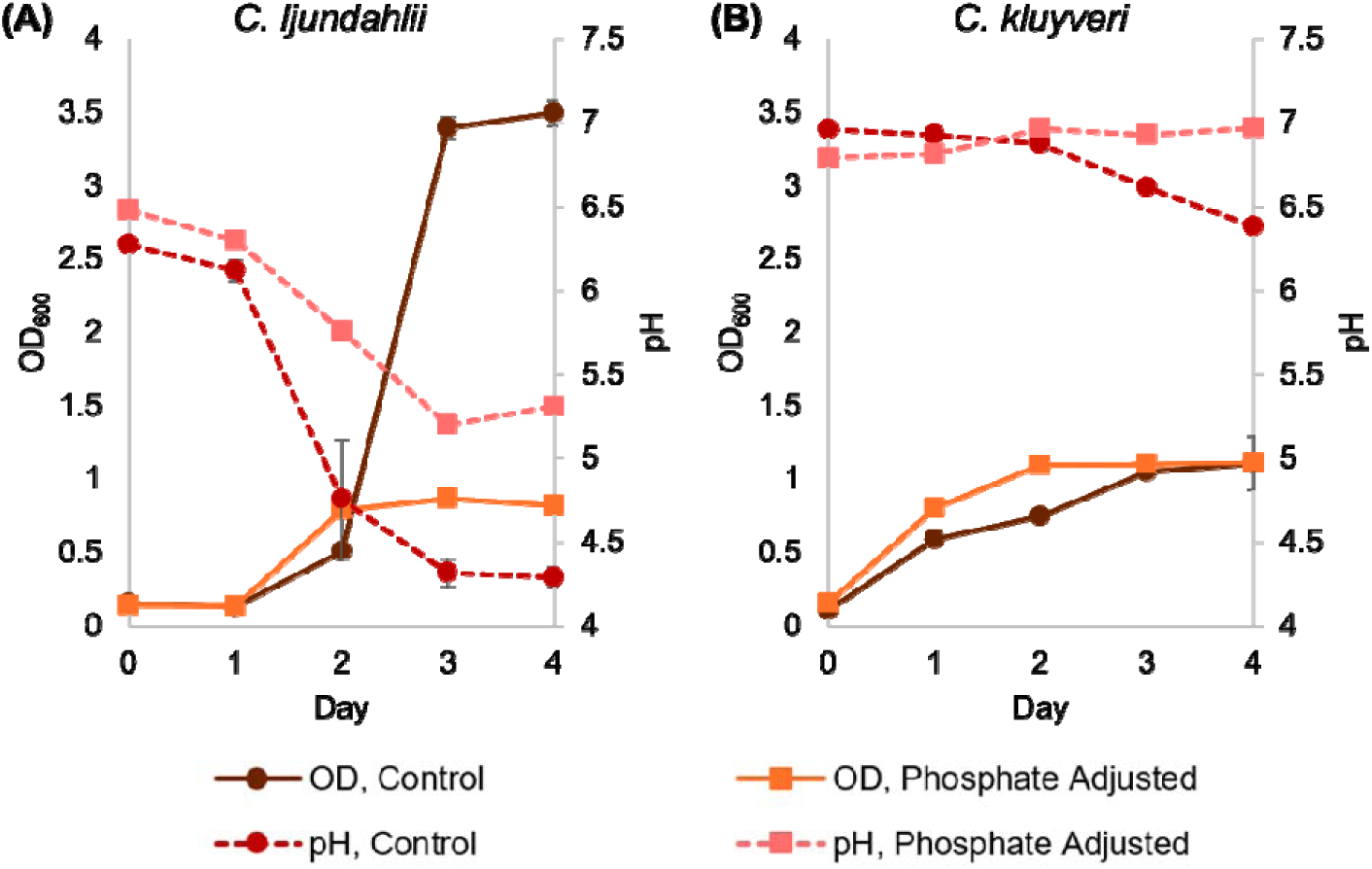
Effects of high phosphate buffering (ca. 140 mM phosphate, Table 1) (“Phosphate Adjusted”) versus control (15 mM phosphate buffer, Table 1) on the biomass formation kinetic profiles (in OD_600_) and pH stability of C. *ljungdahlii* (A) and *C. kluyveri* (B). Two biological replicates were employed per condition.

Taken together, these data support the feasibility of using electrolytes with high-phosphate concentrations without extensive conditioning prior to biological processing. Although elevated phosphate imposes modest growth inhibition on *C. ljungdahlii*, the overall platform remains viable and benefits from enhanced pH stability.

Next, we tested the performance of two stage coculture in converting mixtures of formate, acetate, and ethanol into MCFAs under electrolysis-relevant high-phosphate (about 140 mM) conditions. To evaluate process robustness and kinetic tunability, the coculture was operated with varying initial cell densities (OD = 0.1, 0.3, and 0.5) prior to each stage. Across all conditions, the platform remained operational and achieved efficient chain elongation (Fig. 5). Increasing the initial biomass primarily accelerated the formate-conversion stage: the total process time decreased from 120 h at OD = 0.1 (including 96 hours for formate conversion) to 40 hours at OD = 0.5 (with formate conversion completed within 24 hours). In contrast, the subsequent chain elongation phase was comparatively insensitive to biomass inoculum size (16–24 hours). Final product titers were similar across all groups, consistent with identical substrate inputs (50 mM formate, 40 mM acetate, and 100 mM ethanol). Specifically, butyrate and hexanoate reached ∼23–26 mM and ∼24–25 mM, respectively, regardless of initial OD. These results show that increased initial cell density shortens the formate-to-acetate conversion step, but it does not accelerate the chain elongation step. This suggests that, in our prior experiments, *C. ljungdahlii* biomass was rate-limiting in the first step, whereas *C. kluyveri* biomass was not rate-limiting in the second step.

**Figure 5.**
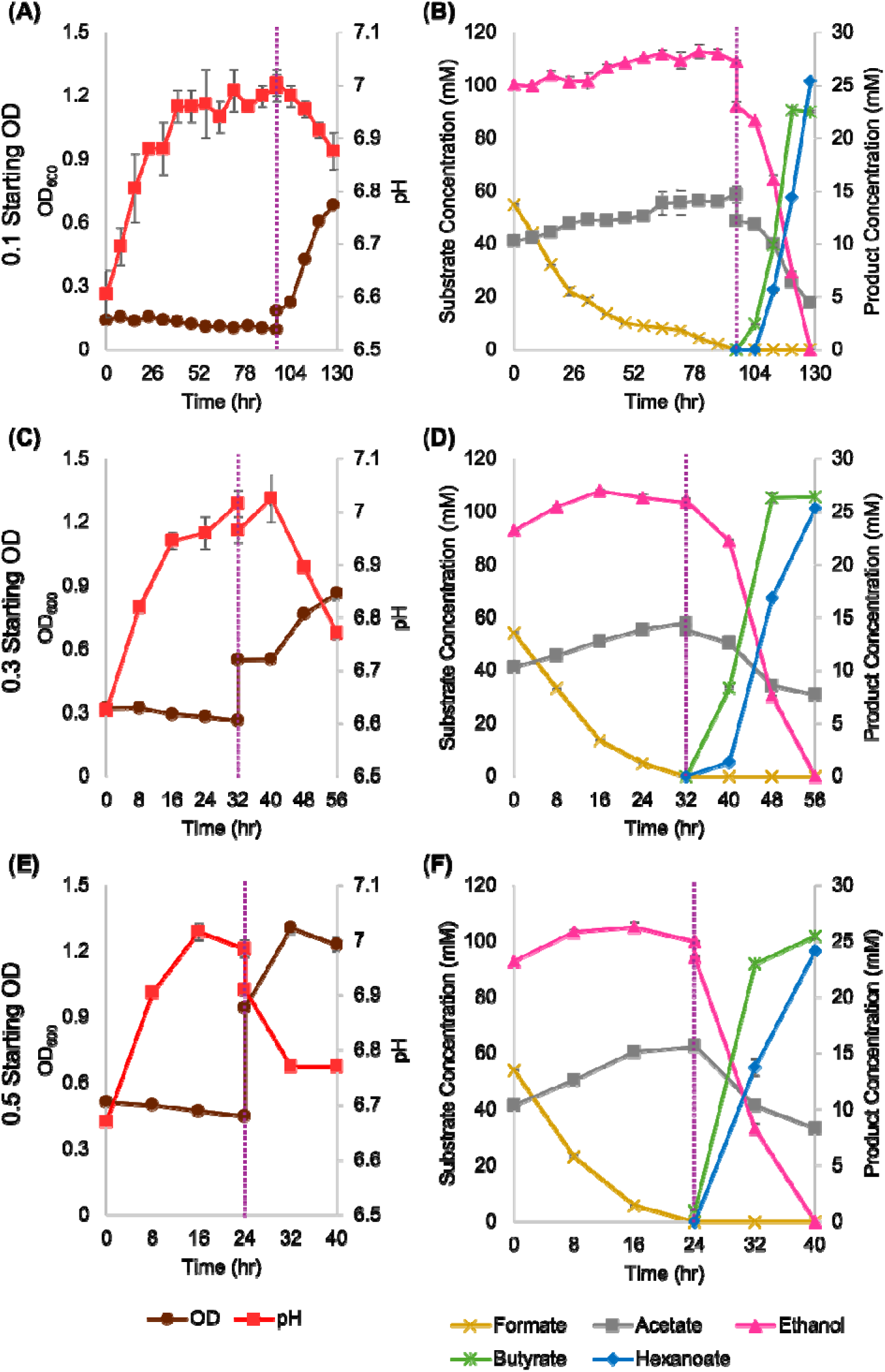
Impact of initial biomass loading on two-stage formate upgrading under high-phosphate (ca. 140 mM) conditions. The dashed vertical lines represent the time of *C. kluyveri* inoculation. (A-B) used a starting OD_600_ of 0.1; (C-D) used a starting OD_600_ of 0.3; (E-F) used a starting OD_600_ of 0.5. (A, C, E) Biomass formation kinetics (in OD_600_) and pH kinetics. (B, D, F) Substrate consumption and product formation kinetics. Two biological replicates were employed per condition.

Next, we tested increased acetate and ethanol concentrations while maintaining constant formate input (50 mM) at an initial OD of 0.5. Elevated acetate/ethanol concentrations prolonged the formate-conversion stage, requiring 88 h for both medium (80 mM acetate/200 mM ethanol) and high (120 mM acetate/300 mM ethanol) conditions (Fig. 6). However, the chain elongation phase remained nearly invariant at ∼16 h, even though substrate concentration was increased, demonstrating that the rate of MCFA synthesis by *C. kluyveri* is fast, efficient, and not limited by *C. kluyveri* cell density at the substrate concentrations tested. MCFA titers scaled proportionally with substrate availability: butyrate increased from 25.4 ± 0.1 mM to 36.2 ± 0.2 mM, and hexanoate from 68.1 ± 0.0 mM to 104.3 ± 0.2 mM. Collectively, these data demonstrate scalable upgrading of electrolysis-derived substrates, including formate, to MCFAs.

**Figure 6.**
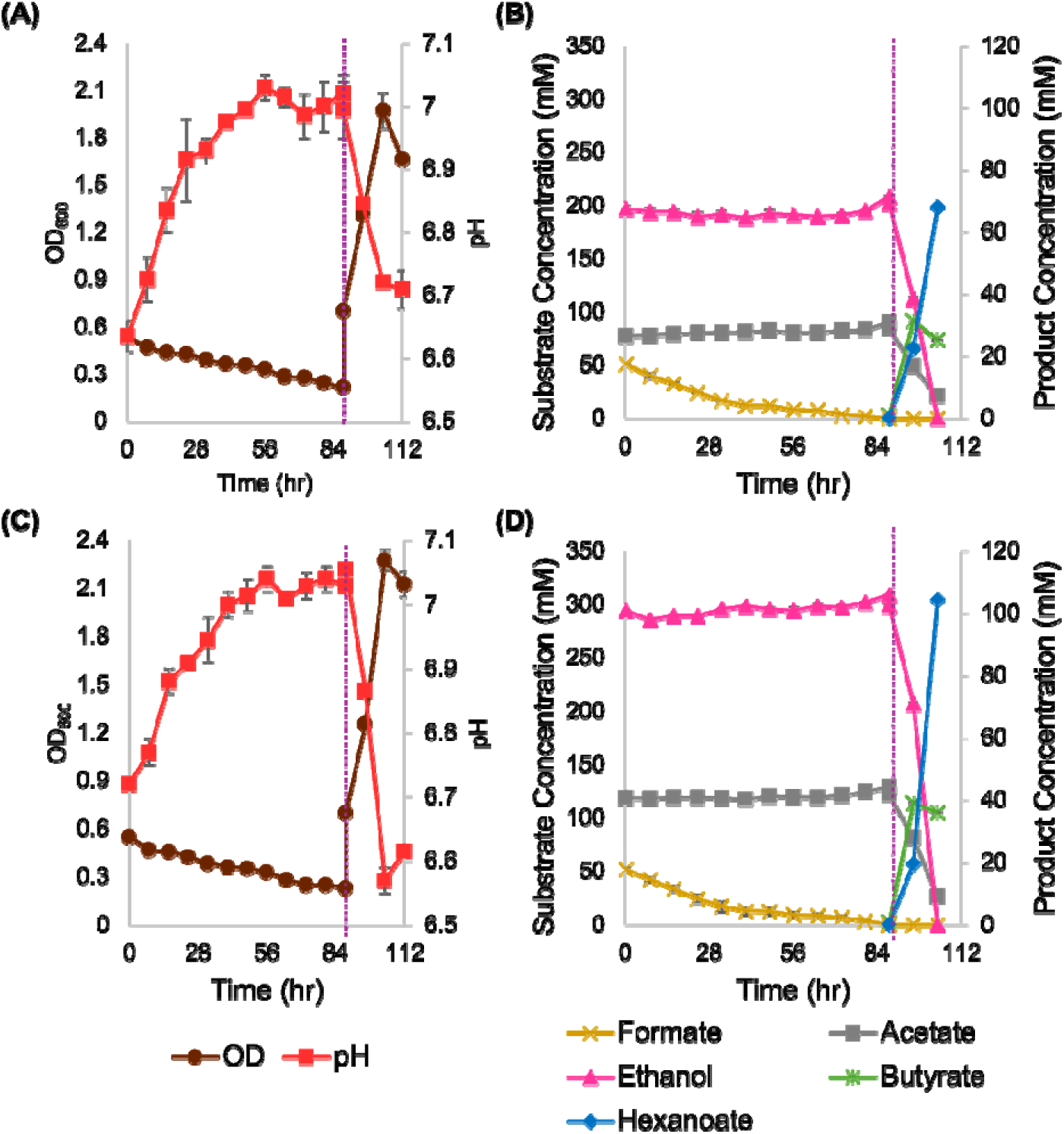
Impact of higher acetate and ethanol loadings on formate upgrading in sequential *C. ljungdahlii* - *C. kluyveri* cocultures under high-phosphate (ca. 140 mM) conditions. (A-B) doubled or (C-D) tripled ethanol and acetate concentrations from those of the experiments of Figure 5. The dashed vertical lines represent the time of *C. kluyveri* inoculation. (A, C) Biomass formation kinetics (in OD_600_) and pH kinetics. (B, D) Substrate consumption and product formation kinetics. Two biological replicates were employed per condition.

### Characterization of *C. ljungdahlii* Biofilms Formation on Glass and Metal Supports

To improve catalyst retention, process stability, and compatibility with a hypothetical continuous reactor configuration, we explored *C. ljungdahlii* biofilms as an immobilized biocatalyst platform for formate conversion. Although *C. ljungdahlii* biofilms have been investigated previously (23), their application as immobilized biocatalysts has not yet been explored. Scanning electron microscopy (SEM) and confocal laser scanning microscopy (CLSM) were used to characterize the morphology, spatial organization, and cellular activity of *C. ljungdahlii* biofilms formed on glass and saddle-shaped metal support materials. Glass support materials were used for ease of analysis. Metal saddle supports were tested because of their biomass compatibility, large surface area for cell attachment, good mechanical strength and light mass. These properties make the metal saddles an excellent potential biofilm support structure and practical packing materials for packed bed reactor configurations (21), which can be used for stable continuous bioprocessing to remove formate prior to chain elongation. SEM imaging revealed dense cell attachment and extensive surface colonization on the metal supports, with continuous biofilm coverage and prominent cell–cell aggregation, consistent with stable establishment of biofilms on the packing material (Fig. 7).

**Figure 7.**
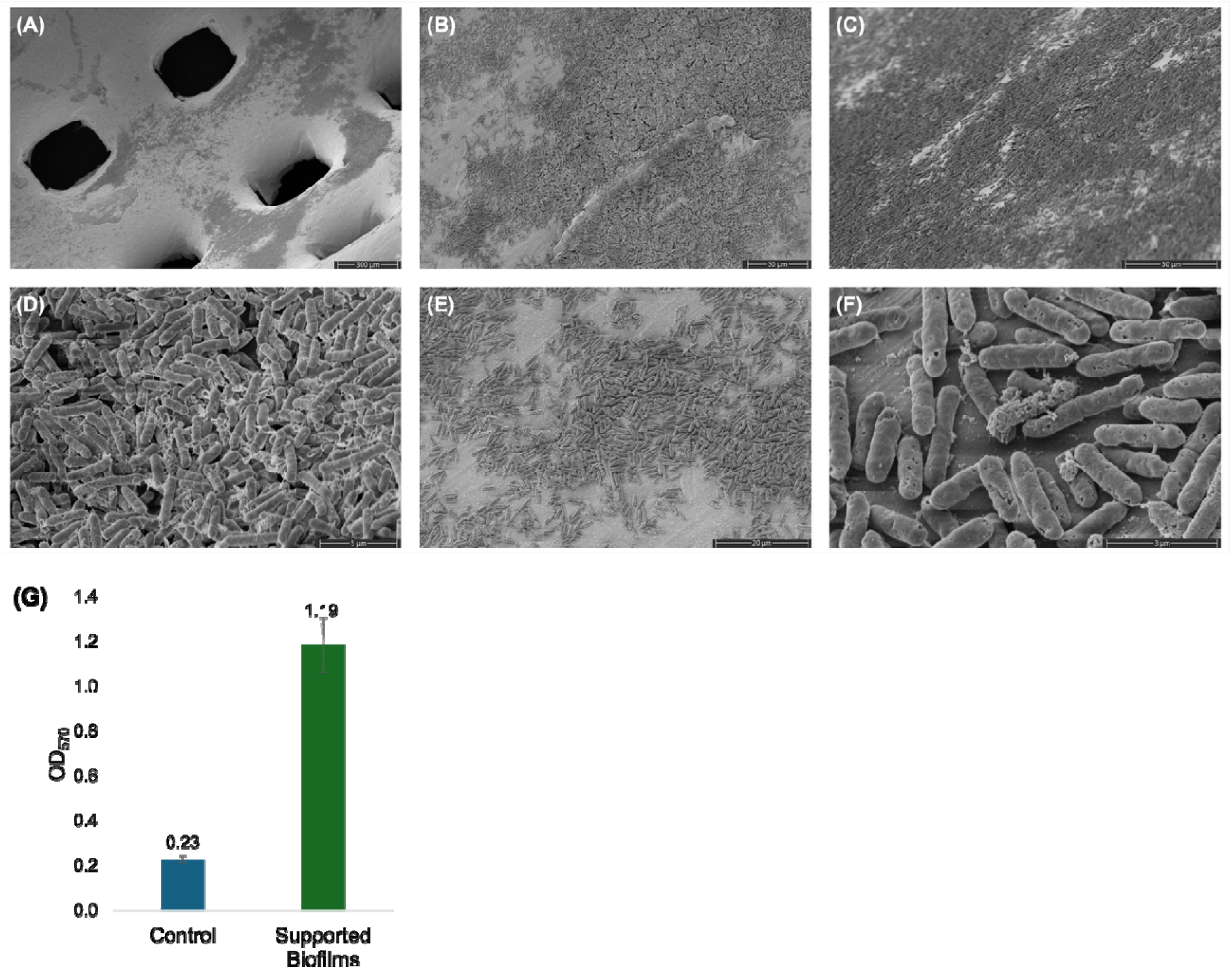
Scanning electron microscopy (SEM) images (A-F) at different magnifications of *C. ljungdahlii* biofilms formed on saddle-shaped metal packing. Scale bars: 300 μm in (A), 30 μm in (B,C), 5 μm in (D), 20 μm in (E), and 3 μm in (F). (G) Quantification of *C. ljungdahlii* biofilm biomass on saddle-shaped metal packing by crystal violet staining (OD_570_).

CLSM coupled with rRNA-targeted fluorescence in situ hybridization (rRNA-FISH) further indicated that many cells within biofilms on both glass and metal exhibited strong rRNA-associated fluorescence signals (Fig. 8), consistent with high active ribosomal content and sustained cellular activity (18).

**Figure 8.**
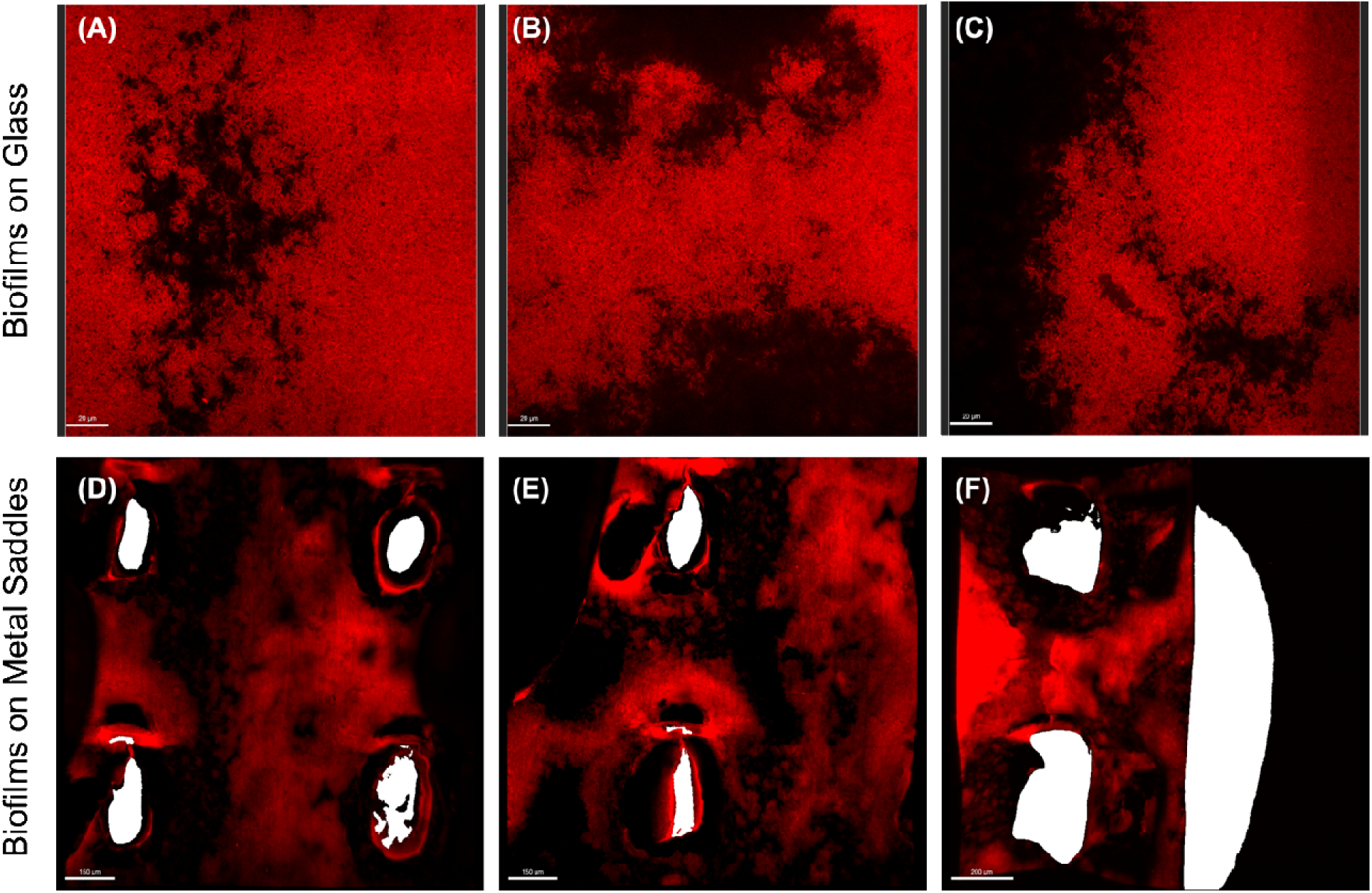
Confocal laser scanning microscopy (CLSM) images of *C. ljungdahlii* biofilms probed by rRNA-targeted fluorescence in situ hybridization (rRNA-FISH). Biofilms formed on glass (A-C) and on saddle-shaped metal supports (D-F). As previously established (see methods), strong rRNA-FISH signals indicate high functional ribosomal content within the biofilm matrix. Scale bars: 20 μm in (A–C), 150 μm in (D, E), and 200 μm in (F).

Together, these results demonstrate that *C. ljungdahlii* can robustly colonize both glass and saddle-shaped metal supports while maintaining high cellular activity within the biofilm matrix—features that support the use of immobilized biofilms for sustained formate conversion in packed-bed or other continuous reactor configurations.

### Formate Conversion by *C. ljungdahlii* Biofilms Supported on Saddle-Shaped Metal Packing

We next quantified formate conversion by the formed biofilms on the metal saddles in the high-phosphate medium, comparing formate removal under an inert N headspace versus a reducing 80:20 (v/v) H /CO headspace across a range of initial formate concentrations. Prior to formate-conversion assays, biofilm loading on the metal saddles was quantified by crystal violet staining (with biomass-free saddles analyzed as an abiotic control). Based on the difference between the *C. ljungdahlii* biofilm-supporting saddles and the control saddles, the average OD of biofilms formed on 40 saddle pieces was ∼0.8 (Fig. 7G), indicating substantial biomass retention on the packing material, but as shown in the rest of the Fig. 7 panels, still only a fraction of the metal surface is covered by the formed biofilm.

Nevertheless, saddle-supported *C. ljungdahlii* biofilms exhibited a dose-dependent formate conversion response under high-phosphate conditions. At 30 mM formate (Fig. 9Aa), *C. ljungdahlii* biofilm removed formate under both N_2_ and H /CO atmospheres. However, H /CO accelerated removal and increased acetate formation. Under N, complete formate removal required ∼4 days and yielded 12.4 ± 0.6 mM acetate (a near stoichiometric conversion; 2 13= 26 mM formate), whereas under 80:20 (v/v) H /CO, formate was depleted within 3 days and acetate formation increased to 28.6 ± 2.1 mM, which corresponds to ∼60 mM of one carbon skeletons, indicating approximately 30 mM CO_2_ fixation based on the WLP.

The benefit of H /CO was most evident at 50 mM formate (Fig. 9B). Under N, biofilms displayed apparent inhibition, consuming only ∼20 mM formate over 5 days with minimal acetate accumulation (thus suggesting that formate was largely converted to CO_2_ from lack of sufficient reducing electrons). In contrast, H /CO alleviated inhibition and enabled complete conversion of 50 mM formate within 3 days, accompanied by the highest acetate titer observed under the tested conditions (∼40 mM, which corresponds to ∼80 mM of C1 skeletons). These results indicate that even with a relative small surface coverage (Fig. 7), biofilm-based formate conversion is effective up to 50 mM formate when sufficient reducing power is supplied. At 70 mM formate, conversion was strongly constrained under both atmospheres and neither condition achieved complete removal (Fig. 9C), indicating that substrate toxicity dominates at elevated formate loads and largely masks the influence of headspace composition.

**Figure 9.**
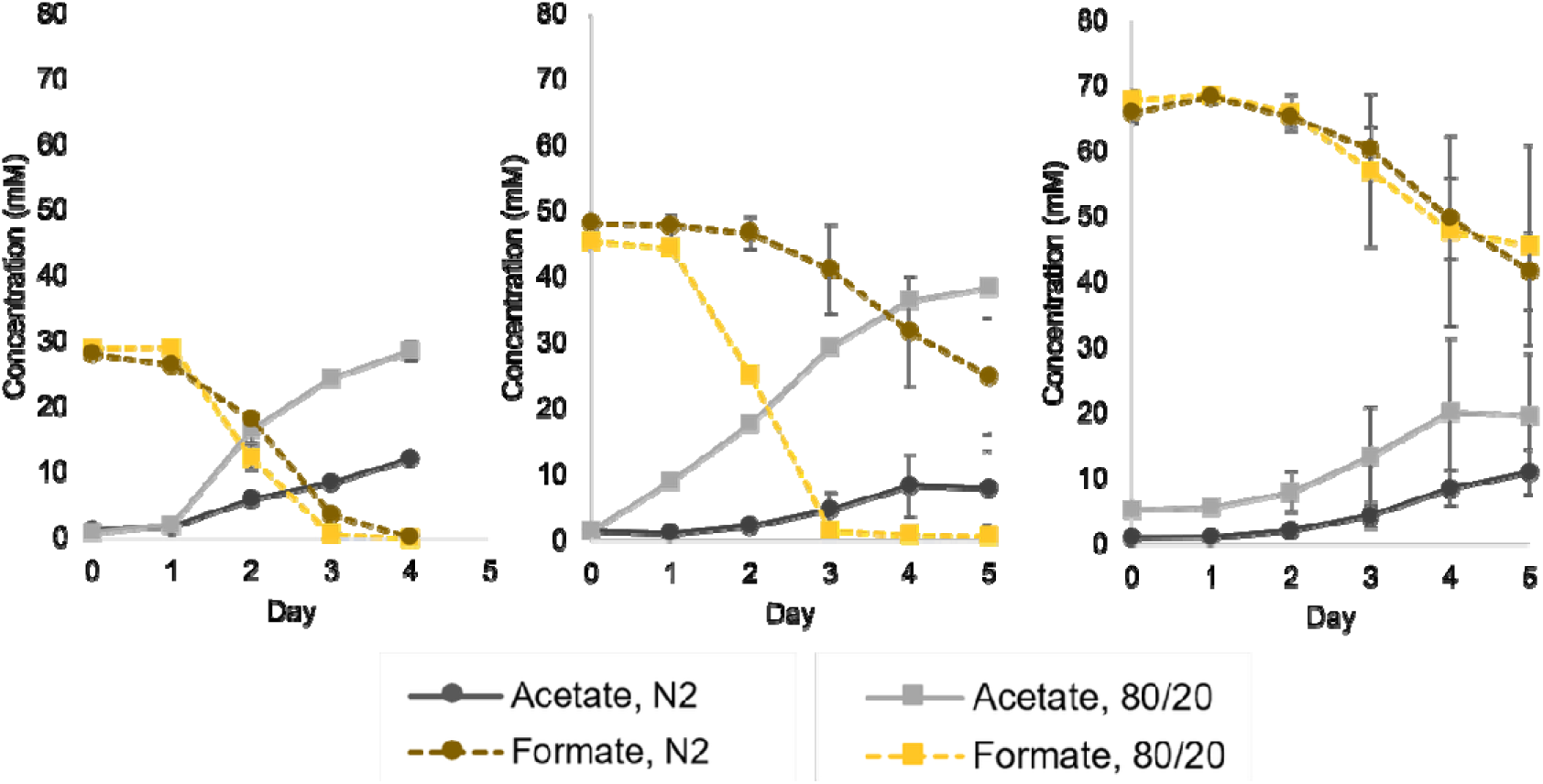
(A-C) Formate assimilation and acetate production during detoxification by *C. ljungdahlii* biofilms on stainless steel saddles under high-phosphate (ca. 140 mM) conditions. Formate loading: 30 mM (A), 50 mM (B), or 70 mM (C), with an atmosphere of N_2_ or 80/20 H_2_/CO_2_.

### Formate Conversion by *C. ljungdahlii* Biofilms in the Presence of Acetate and Ethanol

To better approximate the chemical complexity of CO electrolysis effluents, saddle-supported *C. ljungdahlii* biofilms were challenged with mixed-substrate media containing 40 mM acetate and 100 mM ethanol in addition to formate. Under these conditions, the mixed substrate matrix imposed additional inhibition that reduced formate-conversion performance relative to formate-only assays.At 50 mM, formate conversion under N remained limited and comparable in the presence or absence of acetate/ethanol (Fig. 10A vs. Fig. 9A), consistent with electron limitation and inhibition already dominating under inert conditions. In contrast, under H /CO, the mixed-substrate matrix substantially reduced formate conversion relative to the formate-only condition (Fig. 10A vs. Fig. 9B), indicating that coexisting acetate and ethanol can exacerbate formate stress even when reducing equivalents are available. At 70 mM formate, metabolic activity was strongly suppressed under both gas atmospheres, resulting in only minimal formate conversion (Fig. 10B). This behavior mirrored the formate-only system at the same concentration (Fig. 9C), supporting the conclusion that at elevated formate loads, toxicity becomes the dominant limitation regardless of gas composition.

**Figure 10.**
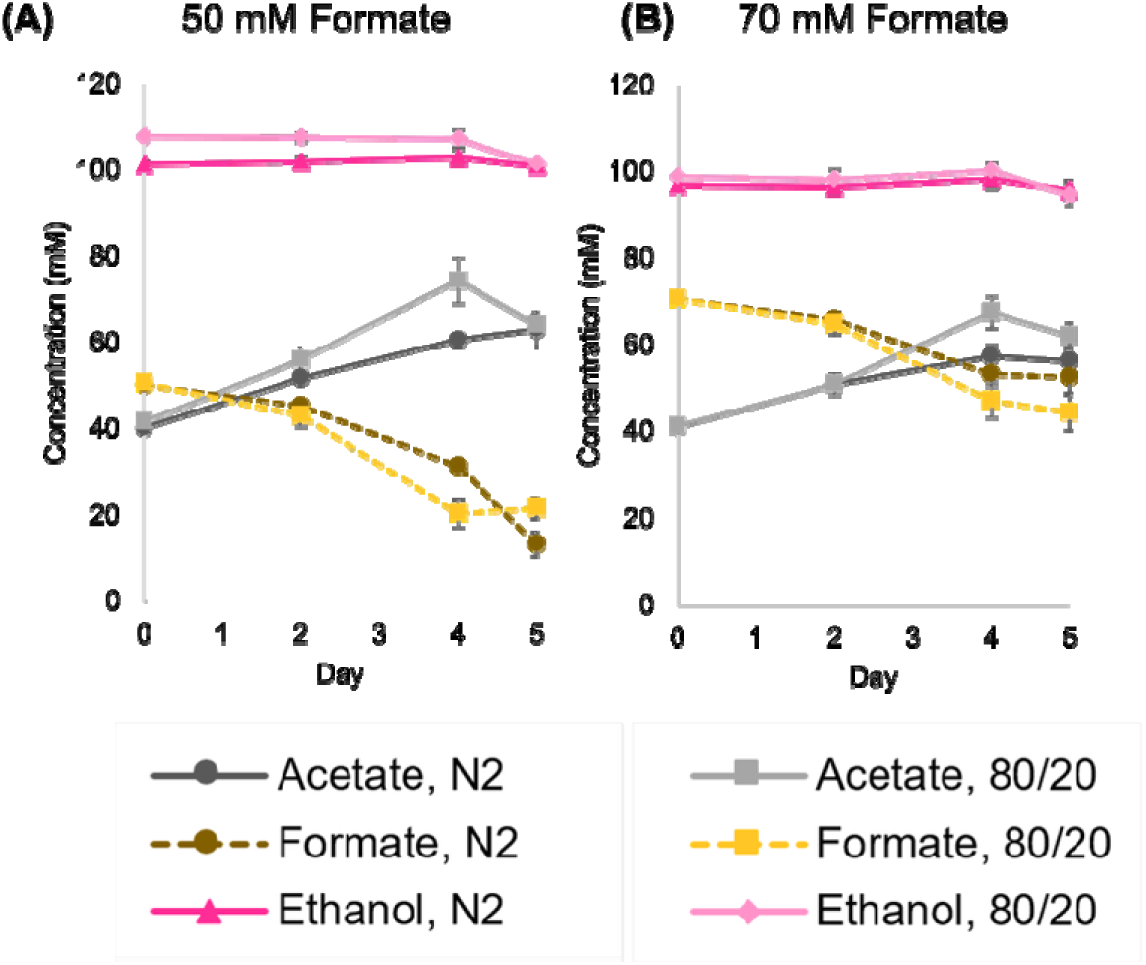
(D-E) Ethanol and acetate formation, and formate uptake kinetics with 50 mM formate (D) or 70 mM formate (E) loading with an atmosphere of N_2_ or 80/20 H_2_/CO_2_. Two biological replicates per condition.

These data support the functional ability of saddle-supported *C. ljungdahlii* biofilms to convert formate from electrocatalytic effluents effluent even with relatively modest biofilm mass (Fig. 9). Noting also the modest size of the employed surface area of the metal saddles, and since formate conversion is proportional to the active mass of *C. ljungdahlii* biofilm, packed bed biofilm bioreactors can be readily developed to handle large volumetric throughputs of formate-containing electrocatalytic effluents.

## CONCLUSIONS & FUTURE DIRECTIONS

This work demonstrated conversion of formate-containing CO RR outputs of ethanol and acetate into MCFAs (butyrate and hexanoate) using *C. ljungdahlii* and *C. kluyveri* via sequential two-step chain elongation. First, we verified that formate did not inhibit either the growth or the chain-elongation activity of *C. kluyveri*. Second, we verified that *C. ljungdahlii* can grow on 50 mM formate with formate as the sole soluble carbon source (in addition to CO_2_ and H_2_ in the headspace). Importantly, *C. ljungdahlii* retained efficient formate consumption even in the presence of acetate and ethanol, supporting its role as a robust substrate-conditioning (“formate conversion”) biocatalyst in mixed-feed environments. Building on this capability, we successfully implemented sequential two-step chain elongation in the *C. ljungdahlii–C. kluyveri* coculture using formate, acetate, and ethanol as mixed substrates, enabling downstream upgrading toward medium-chain fatty acid production. The process remained effective under high-phosphate buffer conditions, demonstrating the versatility and robustness of this coculture strategy in buffered operating windows relevant to practical electrocatalytic applications. Finally, we demonstrated formate conversion by *C. ljungdahlii* biofilms, highlighting the potential of immobilized *C. ljungdahlii* biofilms as biocatalysts for packed-bed reactor configurations. Collectively, these findings provide a foundation for developing scalable, continuous, and mixed-substrate chain-elongation processes for biological upgrading C1-C2 electrocatalytic outputs into higher value MCFAs (C4-C6).

Future research could explore the use of simultaneous (versus sequential) cocultures of *C. ljungdahlii–C. kluyveri,* both in planktonic and biofilm bioreactor settings and combinations of the two types of bioreactors. It would be expected (19) that some or most of the carboxylates produced by *C. kluyveri* would be converted to the corresponding alcohols by *C. ljungdahlii* assuming a good supply of electrons in the form of H_2_. One could also explore the use of other acetogens that might have better tolerance to high formate and phosphate concentrations and/or have broader metabolite capabilities, such as for example *Clostridium carboxidivorans* that produces 4-C chemicals in addition to acetate and ethanol.

An important area of work ought to focus in increasing the concentrations of the CO RR outputs (ethanol and acetate) and the ethanol/acetate ratios to higher than 3 to meet the optimal conditions of *C. kluyveri* chain elongation capabilities. If for example, ethanol concentrations of 600-700 mM can be achieved with ∼ 3:1 ethanol/acetate ratios, then the ability to produce high levels of hexanoate (200-220 mM (10)). pH reduction of the fermentation output might enable substantial reduction of hexanoate’s aqueous solubility to improve separation and overall process economics. Even if ethanol and acetate CO RR outputs do not achieve the >3 ratio, they can still be valorized using supplementation of ethanol from renewable fermentation processes. An appealing concept is to use also the CO_2_ effluents of the ethanol fermentation for CO RR conversion to ethanol and acetate and supplement this output with the ethanol from the fermentation process.

Ethanol and acetate CO RR outputs can be valorized by mixotrophic cultures and cocultures (29, 30) to not only achieve optimal conversion of the CO RR outputs into valuable chemicals, but also expand the metabolite repertoire. For example, as *Anaerotignum neopropionicum* converts ethanol and CO_2_ to propionate, a synthetic co culture of *A. neopropionicum* and *C. kluyveri*, following formate removal by *C. ljungdahlii,* can effectively upgrade dilute streams to odd-chain carboxylic acids (pentanoate and heptanoate) (31).

## Author Contributions

Chao Xu, Conceptualization, Investigation, Methodology, Data curation, Formal analysis, Validation, Writing – original draft, Writing – review and editing.

John D. Hill, Methodology and Investigation. Specifically, John D. Hill assisted with the RNA-FISH experiments and prepared the necessary solutions for these experiments.

Jonathan K. Otten, Visualization, Writing – review and editing. Noah B. Wills, Writing – review and editing.

Eleftherios T. Papoutsakis, Conceptualization, Data curation, Formal analysis, Funding acquisition, Investigation, Methodology, Project administration, Resources, Supervision, Writing – review and editing.

## ACKNOWLEDGEMENTS

This work was supported by the National Science Foundation (NSF) under Award No. 2330245, which funds the Engineering Research Center for Carbon Utilization Redesign through Biomanufacturing-Empowered Decarbonization (CURB).

The authors also thank Dr. Sylvain Le Marchand and Dr. Timothy Chaya at the BioImaging Center, University of Delaware, for their training and assistance with confocal microscopy.

The authors used ChatGPT (OpenAI) as a language-assistance tool to improve clarity and readability of portions of the manuscript. All scientific content, analyses, and conclusions were developed, verified, and approved by the authors.

