## Additional file 1 for "Biological upgrading of C1-C2 products of electrocatalytic CO_2_ reduction to C4-C6 carboxylates"

**Table S1:** Estimated formate concentrations and ratios with ethanol and acetate in relevant electrocatalytic outputs.

| System/Current Density | Main Reported Products | Estimated Formate Concentration | Main Formate/ Ethanol or Acetate Ratio | Notes on Formate | Reference |
| --- | --- | --- | --- | --- | --- |
| Cu, pulsed potential, ~100-200 mA cm^-2^ | Ethanol, CO, ethylene, other C_2_+ compounds | ~10-50 mM | ~3:1 Formate: Ethanol | Formate appears as a minor byproduct in FE plots. Estimated based on typical FE values and product distributions. | [1] |
| Cu with CIBH, >1 A cm^-2^ | Ethanol, ethylene, other C_2_+ compounds | ~5-30 mM | ~2:1 Formate:  Ethanol | Formate suggested to be present. Estimated based on typical FE values and product distributions. | [2] |
| F-doped Cu, 1.6 A cm^-2^ | Ethanol, ethylene | ~25 mM | ~3:4 Formate: Ethanol | Formate appears as a minor byproduct in supplemental FE plots. Estimated based on typical FE values and product distributions. | [3] |
| Ag-modified Cu_2_O, 800 mA cm^-2^ | Ethanol, ethylene | ~25-40 mM | ~3:4 Formate: Ethanol | Formate appears as a minor byproduct in supplemental FE plots. Estimated based on typical FE values and product distributions. | [4] |
| Hydrophobic Cu, ~300 mA cm^-2^ | Ethanol, ethylene | ~5-15 mM | ~2:3 Formate: Ethanol | Formate appears as a minor byproduct in supplemental FE plots. Estimated based on typical FE values and product distributions. | [5] |

| System/Current Density | Main Reported Products | Estimated Formate Concentration | Main Formate/ Ethanol or Acetate Ratio | Notes on Formate | Reference |
| --- | --- | --- | --- | --- | --- |
| MEA, >100 mA cm^-2^ | Ethanol, ethylene, other C_2_+ compounds | <10-20 mM | ~5:8 Formate: Ethanol | Formate not reported as a significant product. Estimated based on typical FE values and product distributions. | [6] |
| N-doped C-capped Cu, 100-300 mA cm^-2^ | Ethanol, ethylene | ~5-10 mM | ~1:2 Formate: Ethanol | Formate appears as a minor byproduct in supplemental FE plots. Estimated based on typical FE values and product distributions. | [7] |
| SnS_2_/Sn_1_-O3G, ~10-30 mA cm^-2^ | Ethanol | ~8-20 mM | ~1:13 Formate: Ethanol | Formate appears as a minor byproduct in FE plots. Estimated based on typical FE values and product distributions. | [8] |
| N-doped nanodiamond, ~1-10 mA cm^-2^ | Acetate | ~10-50 mM | ~2:3 Formate: Acetate | Directly reported | [9] |
| Cu, ~200-1000 mA cm^-2^ | Acetate | <15 mM | ~1:50 Formate: Acetate | Formate not reported as a significant product. Estimated based on typical FE values and product distributions. | [10] |
